# Identification of Monotonically Classifying Pairs of Genes for Ordinal Disease Outcomes

**DOI:** 10.1101/2025.04.10.643941

**Authors:** Océane Fourquet, Daria Afenteva, Kaiyang Zhang, Sampsa Hautaniemi, Martin S. Krejca, Carola Doerr, Benno Schwikowski

## Abstract

In this study, we extend an existing classification method for identifying pairs of genes whose joint expression is associated with binary outcomes to ordinal multi-class outcomes, such as overall survival or disease progression. Our approach is motivated by the need for interpretable classifiers that can provide insights into the underlying biological mechanisms. It can be easily adapted to different research questions, such as identifying gene pair signatures or functional enrichment. We demonstrate that our method is comparable to state-of-the-art classification approaches in terms of performance, while offering the benefit of higher interpretability and adaptability to solve different research questions. Our evaluation on two real-world use cases in glioblastoma and high-grade serous ovarian carcinoma shows that our approach can effectively predict ordinal outcomes and provide novel biological insights. The code is available at https://github.com/oceanefrqt/MBMC.

## 1. Introduction

Advances in transcriptomic profiling, particularly RNA sequencing (RNAseq), have revolutionized the use of gene expression and its role in the study of complex diseases such as cancer. RNAseq provides a comprehensive view of the transcriptome, enabling the identification of differentially expressed genes (DEGs) associated with various cancer-related phenotypes, such as tumor grade, disease progression, or overall survival. Differential expression analysis, where gene expression levels are compared across conditions or groups, is widely used to pinpoint genes that may drive oncogenesis or serve as biomarkers for clinical outcomes. Among DEGs, monotonically differentially expressed genes (MDEGs) (Wang et al. (2015)), whose expression levels consistently increase or decrease across an ordered set of clinical states, such as tumor grades or stages, are of particular interest (Stroggilos et al. (2022)). The identification of MDEGs provides hypotheses about genes that are individually related to cancer progression, offering potential targets for diagnosis and therapy (Tian (2019)).

Although identifying DEGs and MDEGs provides valuable insight into gene expression patterns, these approaches treat genes independently and, therefore, cannot capture higher-order interactions that underlie biological processes. For example, in certain glioblastomas, the interaction between TP53 (a tumor suppressor) and MDM2 (an oncogene that inhibits TP53) is a critical factor for tumor progression. Although TP53 and MDM2 can be identified as DEGs, their individual gene expression does not capture this interaction. To overcome this modeling limitation, approaches such as Top-Scoring Pairs (TSP) (Frank and Hall (2001)) use simple decision rules based on pairs of gene expressions. However, in clinical applications such as predicting tumor grade, disease progression, or patient survival, these outcomes are often on a discrete scale. This makes the task of predicting these outcomes an ordinal classification problem, distinct from the more common categorical classification. Ordinal classification algorithms aim to respect the ordered nature of the outcome, ensuring that predicted categories follow the inherent ranking, making them particularly suited for cancer-related outcomes (Chen (2012)).

In this paper, we provide a new method to construct ordinal multi-class bivariate monotonic classifiers and identify pairs of genes of interest and demonstrate its potential to advance the understanding of the molecular mechanisms underlying cancer.

## 2. Methods

This section describes the data sets we used for our empirical evaluation (Section 2.1), followed by a technical description of the methodologies (Section 2.2). It includes a formal description of the ordinal classification setting (Section 2.2.1), a detailed description of the new MBMC approach (Section 2.2.3) and a comparison with state-of-the-art classification methods (Section 2.2.2). Section 2.3 provides our interpretation of the results.

### 2.1. Data Sets

We chose two data sets with a focus on cancer types with well-defined ordinal outcomes, described in the following sections. After gene filtering, the data sets were split into training (80% of the data) and testing (20% of the data) data sets, using a stratification method that maintains similar ratios between classes in the initial data set and the sub-data-sets.

#### 2.1.1. Glioblastoma Data Set

This data set was obtained by RNAseq from primary tumor samples of 70 patients with glioblastoma in the German Glioma Network. Among the patients are 23 long-term survivors (*>* 36 months overall survival), 16 short-term survivors (*<* 12 months overall survival), and 31 patients with intermediate overall survival, as defined in Reifenberger et al. (2014). Only genes with high variability were selected, using the median absolute deviation technique (Howell (2005)), with a threshold of 0.4, which results in 1,836 genes to analyze (Table 1). Raw RNA sequencing data are publicly available on the NCBI GEO repository (https://www.ncbi.nlm.nih.gov/geo/) with the reference GSE53733. They were generated with the Affymetrix Human Genome U133 Plus 2.0 platform with over 50,000 probesets on chips. Only protein-coding genes were kept, resulting in around 16,000 genes. Multiple probes linked to the same gene were gathered and averaged into a single gene. Gene expression was normalized using the TMM (Trimmed Mean of M-values) method followed by a log1p transformation to stabilize the variance and ensure that the data are on a suitable scale for downstream analysis.

**Table 1.**
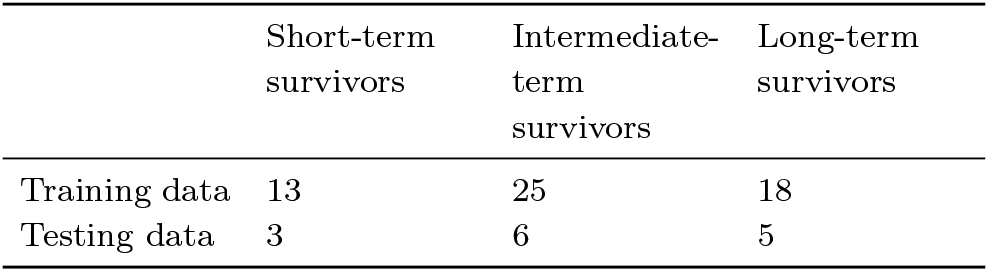
Class balance in glioblastoma training and testing data sets. The numbers represent the number of primary tumor samples from the glioblastoma patients in each category.

|  | Short-term survivors | Intermediate-term survivors | Long-term survivors |
| --- | --- | --- | --- |
| Training data | 13 | 25 | 18 |
| Testing data | 3 | 6 | 5 |

#### 2.1.2. Ovarian High-Grade Serous Carcinoma Data Set

The DECIDER project (https://www.deciderproject.eu) has produced a bulk RNAseq data set of high-grade serous ovarian carcinoma (HGSC) samples, featuring various sample types, including primary tumors, intra-abdominal lesions, and ascites, from a well-characterized cohort (https://clinicaltrials.gov/study/NCT04846933?tab=table). Data were processed as described in Lahtinen et al. (2023). Patients in the cohort underwent surgery, allowing tissue sample collection, followed by platinum-based chemotherapy. Our study focused on the prediction of the time between the last chemotherapy cycle and relapse observation, called the platinum-free interval (PFI), and was based on samples collected before treatment. To account for the heterogeneity of the samples, the PRISM algorithm (Hakkinen et al. (2021)) was used to deconvolute the bulk RNA-seq data into cancer-, immune-, and stromal-specific expression profiles. For this study, only cancer-specific profiles were used in the analysis. To avoid redundancy, the sample with the highest tumor purity score, as estimated by PRISM, was selected for patients with multiple samples. Finally, gene expression data were normalized using the TMM method followed by a log1p transformation. Genes were filtered using the median absolute deviation technique, with thresholds of 1.6, 1.4, and 1.8 for primary tumors, intra-abdominal lesions, and ascites, respectively. The different sub-data-sets are described in Table 2. The classes were determined according to the common clinical classification that considers patients resistant to platinum-based chemotherapy when their PFI is less than 6 months, semi-sensitive when their PFI is between 6 and 12 months, and sensitive when the PFI is greater than one year (Luyckx et al. (2022)).

**Table 2.**
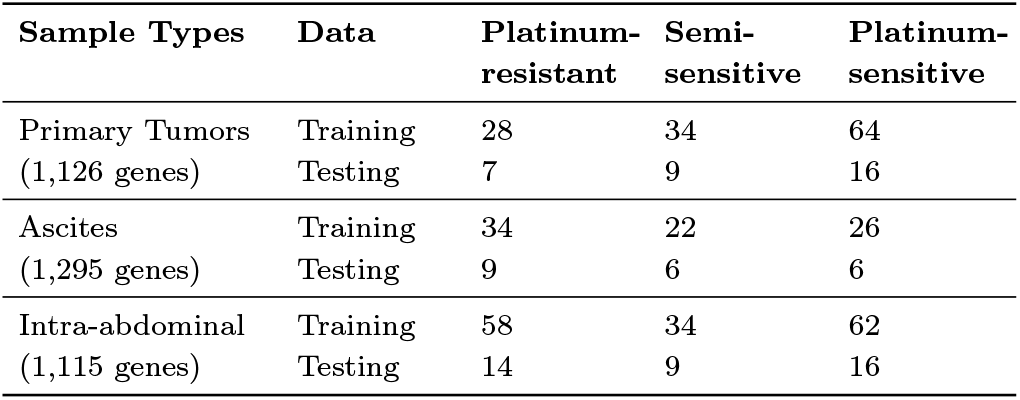
Class balance in training and testing data sets. The numbers represent the number of samples in each category.

| Sample Types | Data | Platinum-resistant | Semi-sensitive | Platinum-sensitive |
| --- | --- | --- | --- | --- |
| Primary Tumors (1,126 genes) | Training | 28 | 34 | 64 |
|  | Testing | 7 | 9 | 16 |
| Ascites (1,295 genes) | Training | 34 | 22 | 26 |
|  | Testing | 9 | 6 | 6 |
| Intra-abdominal (1,115 genes) | Training | 58 | 34 | 62 |
|  | Testing | 14 | 9 | 16 |

### 2.2. Algorithmic Framework

This section defines the concept of ordinal classification (Section 2.2.1), introduces our new approach for multi-class ordinal monotone classification (Section 2.2.3), and summerizes the evaluation metrics as well as the algorithms to which the method is compared (Section 2.2.2).

#### 2.2.1. Ordinal Classification

Ordinal classification (Gutiérrez and García (2016)), corresponds to the task of predicting an outcome with ordered categories such as *low, medium*, and *high*. The key characteristic of it is that it takes into account the **relative ranking** of these categories. For example, we know that *low* is worse than *medium*, and *medium* is worse than *high*.

Monotonic classification is a subtype of the ordinal classification task in which monotonicity constraints are added, meaning that the learned relationship between input features and the outcome is monotonic. A classifier *f* is called **monotonic** if, for any *i* ∈ {1, …, *n*} and any Δ ∈ ℝ, the sign of *f* (…, *x*_*i*_, …) − *f* (…, *x*_*i*_ +Δ, …) does not take on both −1 and 1 over the domain (*x*_1_, …, *x*_*n*_) ∈ ℝ^*n*^.

The performance of ordinal classification can be evaluated according to different metrics. Among the most common ones are (Cardoso and Sousa (2011) and Gaudette and Japkowicz (2009)):

- Accuracy (*Acc*): Ratio between the number of correct predictions and the total number of predictions.
- Accuracy within *n*: Proportion of predictions that are within a certain distance (*n*) of the actual class label.
- Mean Absolute Error (*MAE*): Average difference between the predicted and actual class labels.
- Mean Squared Error (*MSE*): Similar to MAE, but it squares the differences between predicted and actual class labels.
- Cohen’s Kappa (*κ*): statistical measure that evaluates the agreement between predictions actual class labels, correcting for chance agreement.
- Spearman’s Rank Correlation Coefficient: Correlation between the predicted and actual class labels, taking into account the ordinal nature of the classes.
- Matthews Correlation Coefficient (*MCC*): Correlation coefficient using true positives, true negatives, false positives, and false negatives.

All of these metrics either do not take into account the inherent class order or assume that the classes have a fixed, equal spacing between them (Cardoso and Sousa (2011); Gaudette and Japkowicz (2009)). For our study, we have chosen to work with the following metrics: *MAE, Acc, κ*, and *MCC*.

We note that among all the above metrics, *MAE* is one of the most robust common metrics for ordinal classification (Gaudette and Japkowicz (2009)) and fits the flexible frame of ordinal classification. This is why we also use it in our approach. Moreover, we note that the way that the classes are assigned to numbers can have an impact. In order to avoid bias toward some classes, we thus assume in the following use cases that there is an equal absolute distance (of 1) between neighboring classes.

#### 2.2.2. Other Commonly Used Classifiers

The **scikit-learn** (Pedregosa et al. (2011)) library offers numerous implementations of classic classification algorithms, such as **decision trees** (*DT*), **random forests** (*RF*), **logistic regression** (*LR*), **Gaussian processes** (*GP*), and **support vector machines with linear and radial basis function kernels** (*SVM*_linear_ and *SVM*_rbf_). However, in their versions provided by scikit-learn, the algorithms typically treat each class as categorical and independent, thus failing to exploit the information contained in the ordering of the classes (in the case of ordinal classification). This limitation can be bypassed through a simple transformation of the ordinal classification problem into *k* − 1 binary classification problems, where *k* is the number of ordinal classes (Frank and Hall (2001)). In the context of our evaluation, the GridSearch method from scikit-learn was used to systematically explore a predefined range of hyperparameters and identify the best combination of parameters for each model.

#### 2.2.3. Multi-class Bivariate Monotonic Classifier (MBMC)

Bivariate monotonic classifiers (BMC) can be constructed using an efficient dynamic programming-based regression algorithm (Stout (2013)). In the ensembleBMC ensemble classifier, they were used to predict binary disease outcomes from transcriptomic data (Nikolayeva et al. (2018)). We call the extension to the multi-class scenario we present here *Multi-class bivariate monotonic classifier (MBMC)*.

To illustrate the operation of MBMC, let us consider a data set with *n* samples. Each sample belongs to one of *p* ordered classes, with class labels ranging from 1 to *p* (where *p < n*). The goal is to find a set of monotonic functions that separate these ordered classes, minimizing the *L*_1_-error.

MBMC is based on the same regression algorithm as traditional BMC (Stout (2013)), which uses a divide-and-conquer strategy to divide any regression problem with more than two classes into two subproblems when needed, to afterwards merge the resulting solutions. It consists of recursively dividing the classes into subgroups (Figure 1). Initially, all classes are grouped together. We then split them into two subgroups: one containing classes 1 to *k* and the other containing classes (*k*+1) to *p*. Here, *k* is often chosen as the integer part of *p/*2. Stout (2013) proves that the separation function between these two subgroups can be found in *𝒪* (*n* log *n*) time complexity. Following this initial separation, the process can be recursively applied to the resulting subgroups. Each subgroup is further divided into two based on class labels, and a separation function is determined for the new subgroups. This recursive process continues until a separation function is found for each pair of classes. While the complexity of finding a single separation function is *𝒪* (*n* log *n*), the overall complexity of recursively separating all class pairs is likely closer to *𝒪* (*np* log *n*) in the worst case. The approach requires ordered classes.

**Figure 1:**
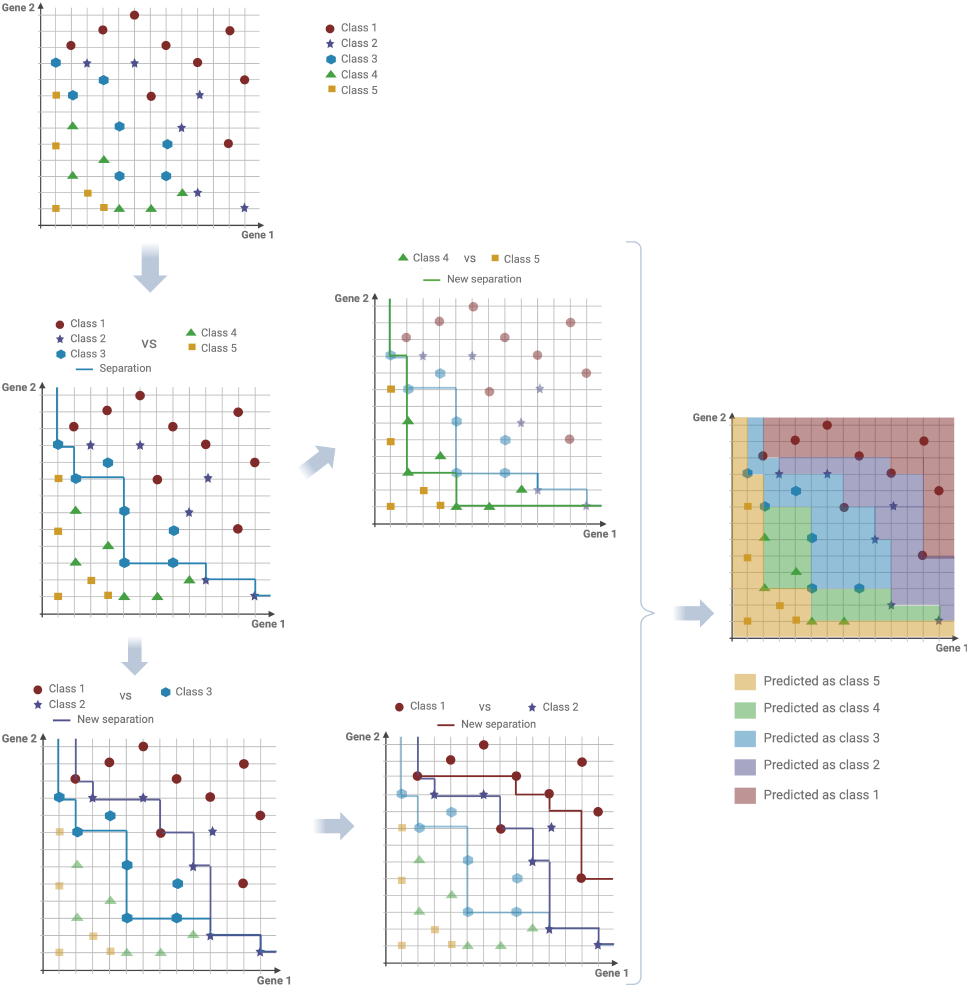
A step-by-step illustration of building a simple MBMC with 5 classes, showcasing the iterative process of creating separation functions and the final classifier, as described in Stout (2013).

Based on the regression algorithm above, MBMC aims to determine the pairs of top-performing gene expressions^1^ through *k* cross-validation. Since a simple brute-force method for calculating the performance of all existing gene pairs in the data set is both time consuming and memory intensive, we instead apply a preselection algorithm that acts as a heuristic to determine well-performing pairs early on (Fourquet et al. (2024)). This preselection identifies and selects the pairs for which the cross-validation is calculated, based on the idea that calculating the MAE on the whole data set (MAE_full_) gives a lower bound on the MAE calculated with cross-validation (MAE_CV_). The algorithm takes as a parameter the minimum number of disjoint pairs, i.e., pairs without genes in common. The details of the algorithm and a mathematical proof of the relationship between both types of MAE are in Section 4.

We note that the number of gene pairs required can vary depending on the specific research question. For instance, identifying a gene signature may only require a small set of 5–10 disjoint pairs, whereas functional enrichment analysis typically benefits from a larger set of 50–200 genes, corresponding to 25–100 disjoint pairs. However, constructing and analyzing gene networks may require a set of more than 100 genes. This flexibility in scale is a key advantage of our approach, as it can be tailored to accommodate different research objectives.

### 2.3. Interpretation and Functional Annotations

The identification of top-performing pairs is interesting from two perspectives: individually and in groups. Individually, each pair puts two genes in relation, which allows capturing their coordinated behavior, such as whether they are both increasing or decreasing in expression and phenotypes, or if their expression patterns are inversely related. MBMCs are designed to strike a balance between simplicity and understandability, ensuring that the identified gene pairs exhibit a robust and easily interpretable monotonic pattern. By avoiding the rigidity of linearity and the opacity of complex relationships, combined with their bi-dimensionality, MBMCs provide a more nuanced and insightful representation of the data, enabling researchers to uncover meaningful and actionable insights that might be obscured by traditional methods.

Furthermore, when considering the top-performing pairs as a group, functional enrichment analyses help uncover pathways that are enriched among these pairs. Functional enrichment analysis was performed using *enrichr* from the **gseapy** library (Zhuoqing Fang (2022)) and the three databases MSigDB Hallmark, MSigDB Oncogenic Signatures, and PID. For each data set, genes after filtering were used as a background for enrichr. The pairs were grouped according to their behavior and the orientation of the relationship. Orientation 1 corresponds to the group of pairs for which both gene expressions increase with the classes. Similarly, Orientation 2 is for pairs whose gene expressions decrease with the classes. And the last group, Orientation 3, is for the mixed signals (one gene increasing and the other decreasing).

## 3. Empirical Evaluation

This section presents an empirical evaluation on the different data sets presented in Section 2.1, including a visualization of the best pairs and a comparison with other methods in terms of performance and interpretability. The best performing MBMCs are selected using the algorithm described in Section 4 according to three distinct scenarios, with the minimum number of disjoint pairs serving as the variable parameter to identify the best performing pairs. This parameter is set to 5, 10, and 20 in each scenario, and they are, respectively, labeled as MBMC-5, MBMC-10, and MBMC-20. Classifiers are trained and tuned on the training data sets, using a 5-fold cross-validation, and each performance is computed on the testing data sets.

### 3.1. Results on Glioblastoma Data Set

Table 3 provides the number of identified top pairs and the number of different genes between these pairs for the three scenarios.

**Table 3.** Summary of the number of top pairs and genes identified in the three scenarios for the glioblastoma data set.

|  | MBMC-5 | MBMC-10 | MBMC-20 |
| --- | --- | --- | --- |
| Number of top pairs | 7 | 23 | 102 |
| Number of genes among the top pairs | 12 | 35 | 112 |

Figure 2 illustrates two of the pairs of genes that perform the best as identified by the MBMC, as well as the associated classifiers, constructed using the competing algorithms (Section 2.2.2). The scattered dots represent the training data points and the colored background corresponds to the classifier built from the data^2^, where each color denotes one of the three classes. These classes are associated with survival outcomes, ranging from the lowest survival rate (Class 0) to the highest survival rate (Class 2), with Class 1 representing an intermediate survival rate. This visualization helps to easily understand the behaviors of the two gene expressions and their relations with the ordinal outcomes.

**Figure 2:**
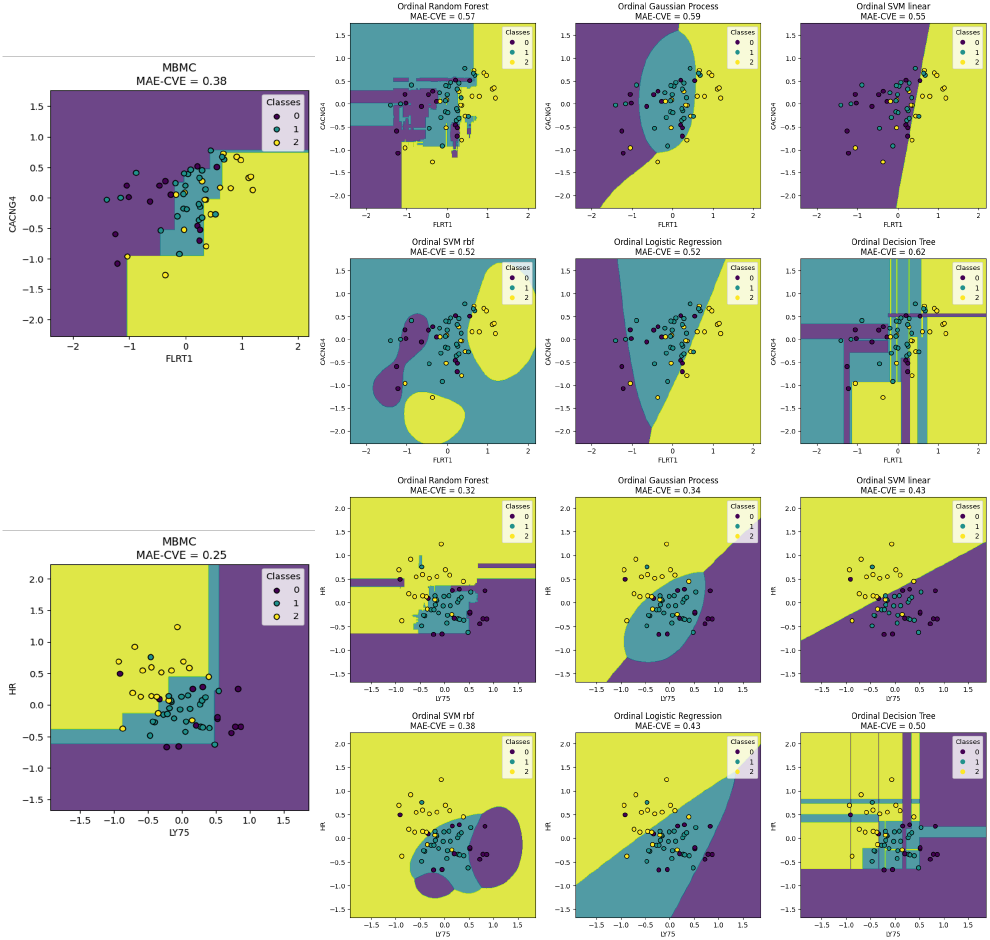
Comparison of the visual representation of the classifiers for two of the best pairs of genes constructed on glioblastoma training data (described in Table 1). The dots represent the training data points and the colored background corresponds to the classifier built from these data points. The two models one on the left are from MBMC, the other models are from the competing approaches (Section 2.2.2).

From a comparable visual perspective, our model stands out for its clear and identifiable pattern, allowing intuitive understanding and generalization of the underlying trend. Although it may not be the most precise, its simplicity enables the formulation of realistic hypotheses that can be applied in real-world scenarios. Notably, our model’s pattern is more generalizable and interpretable than that of a decision tree, which, despite similarities, can be limited by its rigid structure. In contrast, logistic regression, while capable of capturing the monotone trend, often does so with less nuance and detail, failing to provide the same level of insight as our model. However, traditional algorithms are trained on many more than just two genes. A visualization of this order therefore requires a dimensional reduction, such as a PCA, making interpretation all the more difficult, in stark contrast to MBMC.

As the MBMC method produces a set of top-pair classifiers, the performance was calculated for the testing data for each classifier, built with the training data. To obtain a single score per metric for each scenario, the median of the performances across the top pairs was taken. Competing classification algorithms (listed in Section 2.2.2) were fine-tuned with grid search and adjusted to respect ordinal constraints. Since random forest and decision tree algorithms incorporate randomness, we evaluated the median performance of 10 runs. According to the performance ranking (Table 5), the top-performing algorithms for the glioblastoma data set were logistic regression and SVM_rbf_ which achieved the highest rankings in several metrics. In comparison, MBMC algorithms, with the selection parameters set at 5, 10, and 20, did not achieve the highest ranking but still demonstrated competitive performance (Table 4), generally outperforming algorithms such as Gaussian processes and decision trees. Although the performance of MBMC algorithms did not exceed the other top-performing models, they maintained consistent performance across the metrics. In particular, the performance gap between the MBMC algorithms and the models that performed worst, such as Gaussian processes, was substantial, indicating that the MBMC approach remains a viable option for the analysis of the glioblastoma data set.

**Table 4.** Performance evaluation of different models, with metrics including Mean Absolute Error (MAE), Accuracy (Acc), Matthews Correlation Coefficient (MCC), and Cohen’s Kappa for the glioblastoma data set.

|  | Algorithms |  |  |  |  |  |  |  |  |
| --- | --- | --- | --- | --- | --- | --- | --- | --- | --- |
| | MBMC-5 | MBMC-10 | MBMC-20 | RF | DT | LR | GP | $SVM_{linear}$ | $SVM_{rbf}$ |
| MAE | 0.57 | 0.57 | 0.64 | 0.54 | 0.64 | 0.50 | 1.14 | 0.64 | 0.50 |
| MCC | 0.22 | 0.22 | 0.22 | 0.36 | 0.36 | 0.37 | 0.00 | 0.35 | 0.26 |
| $\kappa$ | 0.20 | 0.18 | 0.20 | 0.28 | 0.31 | 0.29 | 0.00 | 0.34 | 0.13 |
| Acc | 0.50 | 0.50 | 0.50 | 0.57 | 0.54 | 0.57 | 0.21 | 0.57 | 0.50 |

**Table 5.** Ranking of models based on their performance, with the top-performing model ranked 1st and subsequent models ranked accordingly, applied to the glioblastoma data set.

|  | Algorithms |  |  |  |  |  |  |  |  |
| --- | --- | --- | --- | --- | --- | --- | --- | --- | --- |
| | MBMC-5 | MBMC-10 | MBMC-20 | RF | DT | LR | GP | $SVM_{linear}$ | $SVM_{rbf}$ |
| MAE | 4 | 4 | 6 | 3 | 6 | 1 | 9 | 6 | 1 |
| MCC | 7 | 7 | 6 | 2 | 3 | 1 | 9 | 4 | 5 |
| $\kappa$ | 5 | 7 | 5 | 4 | 2 | 3 | 9 | 1 | 8 |
| Acc | 5 | 5 | 5 | 1 | 4 | 1 | 9 | 1 | 5 |

Among the selected pairs, the identification of a signature of gene pairs and a corresponding **ensemble model** offers a promising approach to improve predictive performance. By focusing on a compact signature, ideally comprising around 10 genes or fewer, we can use the results of the first scenario. To construct our ensemble classifier, we selected five non-redundant gene pairs that do not share common genes (Figure 3). We then trained this ensemble classifier on the training data, where each pair as an MBMC contributed to a majority vote. Upon testing this ensemble classifier on the testing data, we achieved a **MAE of 0.46**, which surpasses the performance of all other methods explored. This result shows that multiple high-performing gene pairs can be combined into strong predictors.

**Figure 3:**
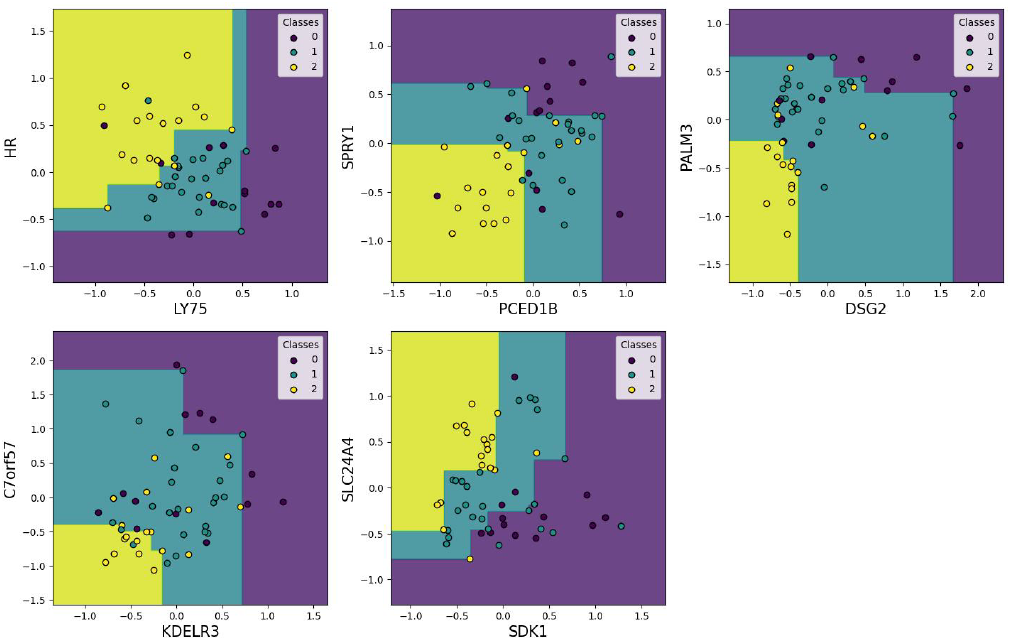
Five non-redundant gene pairs chosen for the MBMC ensemble model, predicting overall survival for glioblastoma patients.

After pathway enrichment analysis, it was determined that the pairs of genes in Orientation 1 were enriched for pathways that are typically associated with breast cancer, including early estrogen response, SRC UP.V1 DN, LTE2 UP.V1 DN, and EGFR UP.V1 DN. It is possible that the underlying biological processes regulated by these pathways, such as cell proliferation and survival, are also relevant to glioblastoma. For example, the SRC pathway is known to play a role in cell migration and invasion, which are also hallmarks of glioblastoma. Similarly, the EGFR pathway is often dysregulated in glioblastoma, leading to increased cell proliferation and survival. Orientation 2 pairs were enriched for the epithelial-mesenchymal transition (EMT) and PDGF UP.V1 DN. The EMT pathway is a process linked to tumor progression and invasiveness, which correlates with a poorer prognosis in glioblastoma. The PDGF UP.V1 DN pathway involves genes down-regulated in neuroblastoma cells in response to Platelet-Derived Growth Factor (PDGF) stimulation. The last group of gene pairs is enriched for SNF5 DN.V1 DN, ATF2 UP.V1 DN, BMI1 DN MEL18 DN.V1 DN, and RELA DN.V1 UP. These pathways involve genes down-regulated in response to perturbations, such as the knockout of SNF5, a tumor suppressor gene, or the over-expression of ATF2, a transcription factor involved in cell growth and survival. The BMI1 DN MEL18 DN.V1 DN pathway, which is associated with the down-regulation of genes involved in stem cell self-renewal, is also notable, as it suggests a potential link between glioblastoma and cancer stem cell biology. The RELA DN.V1 UP pathway, which involves genes up-regulated after the knockdown of the NF-*κ*B subunit RELA, may indicate a role for inflammatory signaling in glioblastoma.

### 3.2 Results on Ovarian High-Grade Serous Carcinoma

The first section compares the results between subtypes, while the second compares them with other classification methods.

#### 3.2.1. Differences Between Sample Types

The results of the three scenarios, including the number of pairs obtained and the number of distinct genes that comprise them, are summarized in Table 6. For the three scenarios, the intra-abdominal data set is the one for which a higher number of top pairs is identified. The other two data sets are quite similar in terms of numbers.

**Table 6.** Summary of the number of top pairs and genes identified across the different sample types in the three scenarios.

|  |  | Primary Sites | Ascites | Intra-Abdominal |
| --- | --- | --- | --- | --- |
| Number of top pairs | MBMC-5 | 12 | 12 | 23 |
|  | MBMC-10 | 24 | 37 | 58 |
|  | MBMC-20 | 102 | 81 | 198 |
| Number of genes among the top pairs | MBMC-5 | 20 | 17 | 26 |
|  | MBMC-10 | 36 | 50 | 63 |
|  | MBMC-20 | 115 | 98 | 175 |

A statistical test was performed to determine whether the observed overlap of genes between two top pairs groups (Table 7) was significantly greater than what would be expected by random chance. The overlap of genes Top pairs from primary sites and from intra-abdominal sites indeed overlapped significantly more than by pure chance (p-value of 2e−5), but not in the two other site comparisons. Interesting, ascites also distinguish themselves biologically from the two other tissue sites, through their character as a *fluid* tissue, and its specific oncological classification of a distant metastasis.

**Table 7.** Overlapping genes between the top pairs in the subtypes.

|  | Primary Sites | Ascites | Intra-Abdominal |
| --- | --- | --- | --- |
| Primary Sites | 115 | 5 | 21 |
| Ascites |  | 98 | 9 |
| Intra-Abdominal |  |  | 175 |

To gain a better understanding of the performance of the MBMC algorithms across different gene pairs and subtypes of samples, the MAE values were calculated for all possible genes. pairs. Figure 4 visualize the distribution of performance. The MAE distributions of every pair for the three subtypes of samples were close to normal distributions. The distribution of MAE values for ascites was more spread out compared to the other two subtypes, suggesting that there was more variability. The distribution of MAE values for intra-abdominal metastasis was more peaked and has a shorter tail, indicating that the MAE values was more concentrated around a central value. The distribution of MAE values for primary sites was similar to that of intra-abdominal metastasis, but skewed toward lower MAE values.

**Figure 4:**
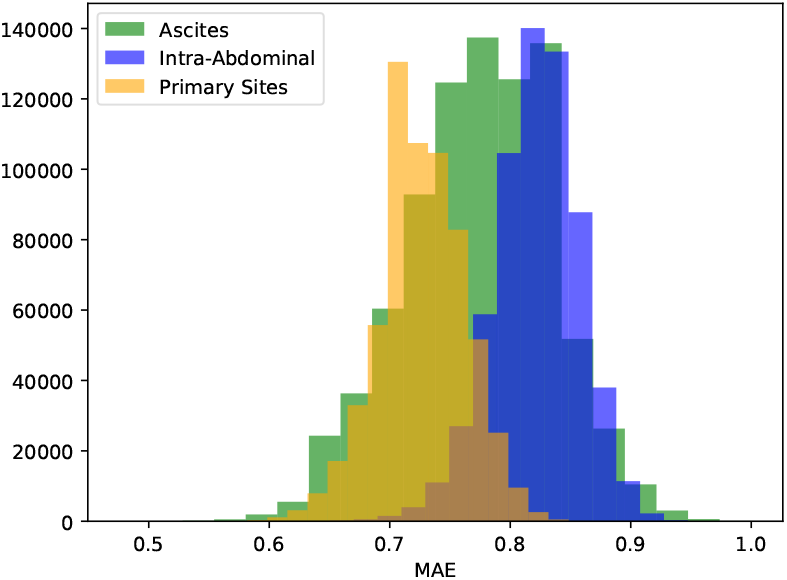
Distribution of mean absolute error values for all possible gene pairs across different subtypes of samples.

To assess the significance of the observed MAE value, we determined whether the MAE of the top pairs was significantly better than what would be expected by chance. First, the class labels were randomly permuted. Then, 1,000 gene pairs were randomly selected, and their MAE values computed. The frequency with which the randomized MAE value was less than or equal to the observed MAE value is used as a *p*-value. For the three data sets, this test results in *p*-values lower than 0.05, implying statistical significance at this level.

#### 3.2.2. Performance Comparison with Other Algorithms to Predict PFI

We compared our MBMC method with standard classification algorithms on the three data sets. Table 8 shows the actual values of the metrics used to evaluate the performance of the algorithms, while Table 9 shows the rank of the algorithms according to their performance.

**Table 8.**
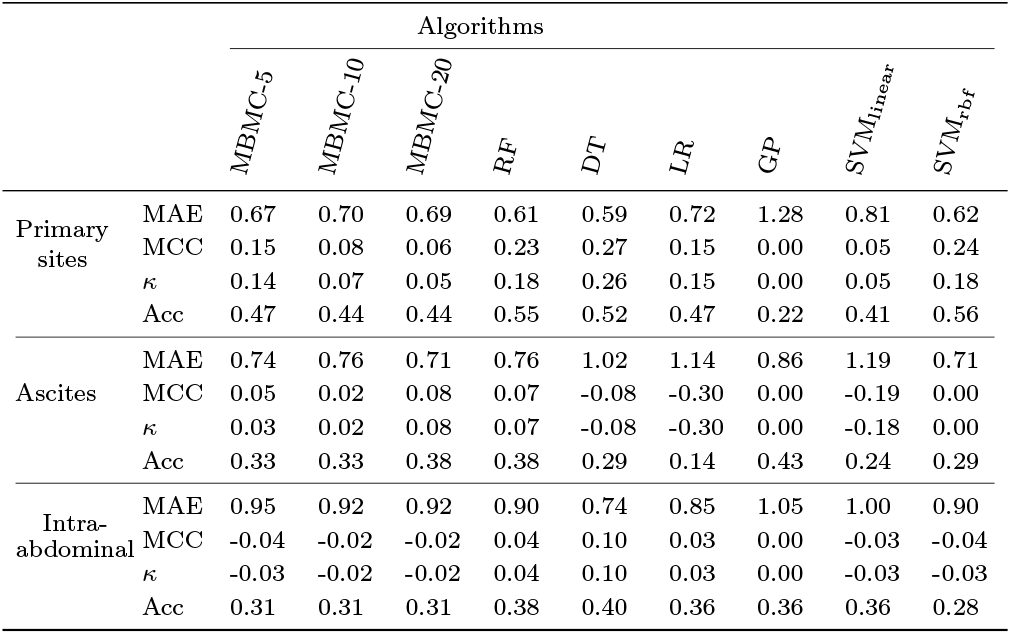
Performance evaluation of different models, with metrics including Mean Absolute Error (MAE), Accuracy (Acc), Matthews Correlation Coef- ficient (MCC), and Cohen’s Kappa, accross the three subtypes of samples.

|  |  | Algorithms |  |  |  |  |  |  |  |  |
| --- | --- | --- | --- | --- | --- | --- | --- | --- | --- | --- |
|  |  | MBMC-5 | MBMC-10 | MBMC-20 | RF | DT | LR | GP | SVM<sub>linear</sub> | SVM<sub>rbf</sub> |
| Primary sites | MAE | 0.67 | 0.70 | 0.69 | 0.61 | 0.59 | 0.72 | 1.28 | 0.81 | 0.62 |
|  | MCC | 0.15 | 0.08 | 0.06 | 0.23 | 0.27 | 0.15 | 0.00 | 0.05 | 0.24 |
| | $\kappa$ | 0.14 | 0.07 | 0.05 | 0.18 | 0.26 | 0.15 | 0.00 | 0.05 | 0.18 |
|  | Acc | 0.47 | 0.44 | 0.44 | 0.55 | 0.52 | 0.47 | 0.22 | 0.41 | 0.56 |
| Ascites | MAE | 0.74 | 0.76 | 0.71 | 0.76 | 1.02 | 1.14 | 0.86 | 1.19 | 0.71 |
|  | MCC | 0.05 | 0.02 | 0.08 | 0.07 | -0.08 | -0.30 | 0.00 | -0.19 | 0.00 |
| | $\kappa$ | 0.03 | 0.02 | 0.08 | 0.07 | -0.08 | -0.30 | 0.00 | -0.18 | 0.00 |
|  | Acc | 0.33 | 0.33 | 0.38 | 0.38 | 0.29 | 0.14 | 0.43 | 0.24 | 0.29 |
| Intra-abdominal | MAE | 0.95 | 0.92 | 0.92 | 0.90 | 0.74 | 0.85 | 1.05 | 1.00 | 0.90 |
|  | MCC | -0.04 | -0.02 | -0.02 | 0.04 | 0.10 | 0.03 | 0.00 | -0.03 | -0.04 |
| | $\kappa$ | -0.03 | -0.02 | -0.02 | 0.04 | 0.10 | 0.03 | 0.00 | -0.03 | -0.03 |
|  | Acc | 0.31 | 0.31 | 0.31 | 0.38 | 0.40 | 0.36 | 0.36 | 0.36 | 0.28 |

**Table 9.**
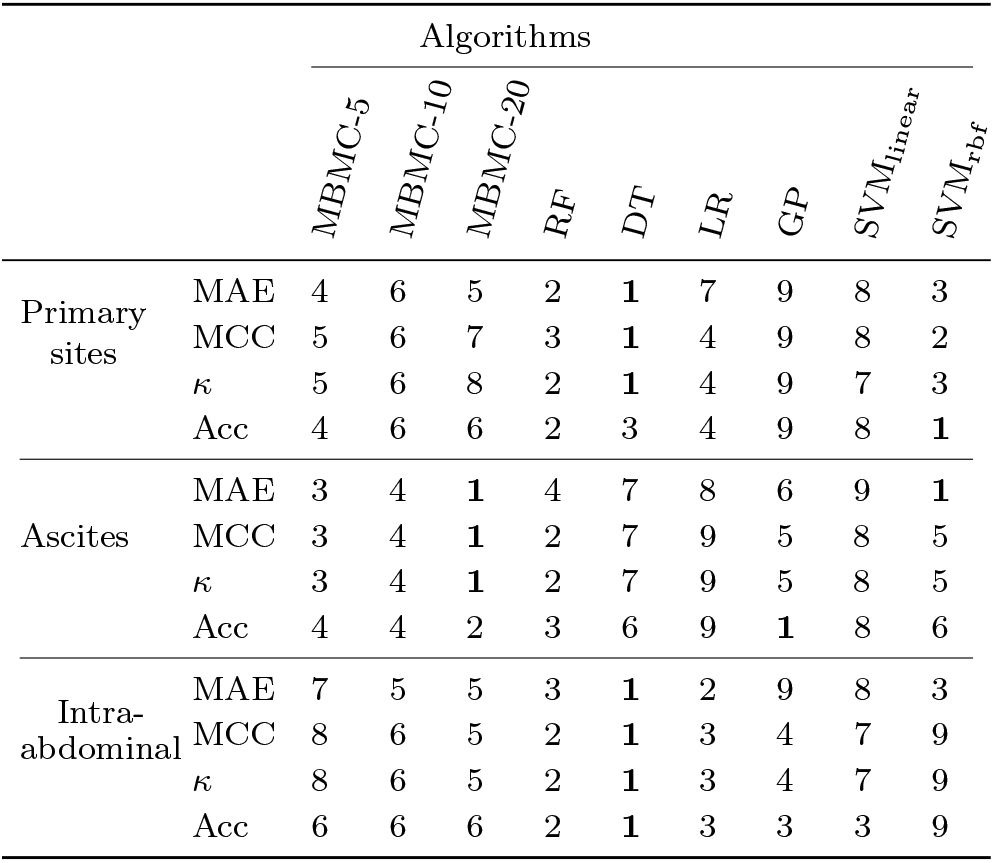
Ranking of models based on their performance, with the top-performing model ranked 1st and subsequent models ranked accordingly, accross the three subtypes of samples.

|  |  | Algorithms |  |  |  |  |  |  |  |  |
| --- | --- | --- | --- | --- | --- | --- | --- | --- | --- | --- |
|  |  | MBMC-5 | MBMC-10 | MBMC-20 | RF | DT | LR | GP | SVM<sub>linear</sub> | SVM<sub>rbf</sub> |
| Primary sites | MAE | 4 | 6 | 5 | 2 | 1 | 7 | 9 | 8 | 3 |
|  | MCC | 5 | 6 | 7 | 3 | 1 | 4 | 9 | 8 | 2 |
| | $\kappa$ | 5 | 6 | 8 | 2 | 1 | 4 | 9 | 7 | 3 |
|  | Acc | 4 | 6 | 6 | 2 | 3 | 4 | 9 | 8 | 1 |
| Ascites | MAE | 3 | 4 | 1 | 4 | 7 | 8 | 6 | 9 | 1 |
|  | MCC | 3 | 4 | 1 | 2 | 7 | 9 | 5 | 8 | 5 |
| | $\kappa$ | 3 | 4 | 1 | 2 | 7 | 9 | 5 | 8 | 5 |
|  | Acc | 4 | 4 | 2 | 3 | 6 | 9 | 1 | 8 | 6 |
| Intra-abdominal | MAE | 7 | 5 | 5 | 3 | 1 | 2 | 9 | 8 | 3 |
|  | MCC | 8 | 6 | 5 | 2 | 1 | 3 | 4 | 7 | 9 |
| | $\kappa$ | 8 | 6 | 5 | 2 | 1 | 3 | 4 | 7 | 9 |
|  | Acc | 6 | 6 | 6 | 2 | 1 | 3 | 3 | 3 | 9 |

Analyzing both tables, we observe the following results:

- **Primary sites**: With an MAE of 0.59, the decision tree demonstrated a strong ability to predict PFI classes. Although the MBMC algorithms did not quite match the performance of the decision tree, they still showed competitive results, with MAE values ranging from 0.67 to 0.70. In terms of accuracy, the trend was similar. The performance of the MBMC algorithms was significantly better than that of other models, such as Gaussian Process and SVM_linear_, which had much higher MAE values. Overall, while MBMC algorithms may not be the best choice in terms of performance, they are still a viable option and can provide competitive results.
- **Ascites**: The MBMC-20 algorithm stood out as the best performer, achieving either the best or second-best performance among all models. Its MAE value was comparable to that of a random forest. Although MBMC-20 did not significantly outperform the other models, it was still a notable achievement. Furthermore, the other MBMC algorithms, MBMC-5 and MBMC-10, also demonstrated relatively strong performance, with MAE values that were comparable to those of other models, excluding outliers such as decision tree, logistic regression and SVM_linear_, which exhibited high MAE values. In general, MBMC algorithms showed promising results in predicting ascites.
- **Intra-abdominal metastasis**: The decision tree algorithm generally emerged as the top performer, closely followed by random forest and logistic regression. In contrast, the MBMC algorithms fell slightly behind the top performers, with MAE values that were somewhat higher than those of DT, RF, and LR. However, it should be noted that the MBMC algorithms were still not among the worst performers, and their results were more comparable to those of Gaussian Process and SVM_linear_. Overall, while the MBMC algorithms may not have been the best choice for predicting intra-abdominal metastasis, they still showed reasonable performance and were not significantly outperformed by all other models.

For the data sets of three subtypes, enrichment analysis did not yield significant results, suggesting that identified gene pairs may not be sufficient to uncover the underlying biological mechanisms. A way to overcome this could be to consider a larger number of gene pairs. Moreover, there might be other possible reasons for this inconclusive enrichment analysis. One possibility is that higher-order interactions between genes are at play, which cannot be captured with only two genes. It is also possible that the identified gene pairs are involved in unknown or uncharacterized pathways that are not represented in the used databases. The relatively small number of patients in the study may also contribute to the lack of significant findings, as larger sample sizes are often required to detect subtle but significant differences. Finally, MBMCs enforce monotonicity relationships between genes and the PFI, which is a big assumption about the relationship.

Although pathway enrichment analysis had not produced statistically significant results in our study, examining individual gene pairs may still provide valuable insights (Figure 5).

**Figure 5:**
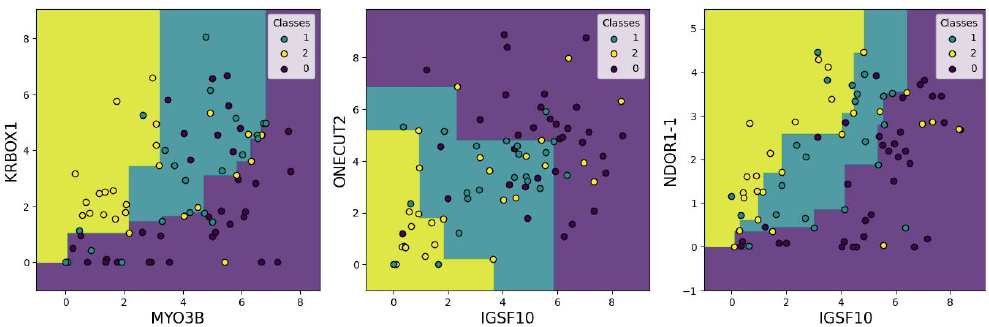
Visualization of three of the top-performing gene pairs, with the MBMC constructed on training data of ascites samples.

## 4. Technical Aspects of the Identification of the Top-Performing Gene Pairs

This section describes the algorithm with which the best MBMCs are identified and the mathematical property that allows for fast preselection. It is a generalization of a preselection algorithm for BMCs (Fourquet et al. (2024)).

### 4.1. Theoretical Property

To introduce the preselection algorithm, we require a few definitions: Let *S* be a non-empty set of data points, and let *P* be a partition of *S*. Let *C* be a monotonic classifier over *S* and *C*_*S*_ an *L*1-optimal monotonic classifier over *S*.

For any set *S*^′^, for any partition *P* ^′^ of *S*^′^, and for any monotonic classifier *C*^′^, let *E*(*C*^′^, *S*^′^) denote the *L*1-error (*L*_1_) of *C*^′^ over *S*^′^, and let *E*(*C*^′^, *P* ^′^) denote the *L*_1_ of *C*^′^ over *P* ^′^. Then, 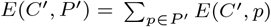 and 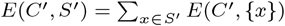.

Moreover, the *L*_1_ of *C*^′^ over *S*^′^ can be broken down into the sum of the *L*_1_ of *C*^′^ over each element *x* in *S*^′^. The elements in *S*^′^ can be grouped into partitions *p*, so the *L*_1_ of *C*^′^ over *S*^′^ is also equal to the sum over all parts *p*in *P* ^′^, and then summing over each element *x* within each partition. It can be simplified to the sum of the *L*_1_ of *C*^′^ over all partitions *p*. Ultimately, this means that the *L*_1_ of *C*^′^ over *S*^′^ is equal to the *L*_1_ of *C*^′^ over *P* ^′^.

#### Theorem 4.1.

*For any non-empty set S and for any p* ∈ *P, it holds that* 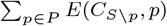.

*Proof* 1.For all parts *p* in *P*, the *L*_1_ of *C*_*S*_ over *S* is lower than or equal to the *L*_1_ of *C*_*S*\*p*_ over the subset *S*. This is because *C*_*S*_ is the optimal monotonic classifier over *S* in terms of *L*1−optimality.

2. For all parts *p* in *P*, the *L*_1_ of *C*_*S*_ over *S* \ *p* is greater than or equal to the *L*_1_ of *C*_*S*\*p*_ over the subset *S* \ *p*. This is because *C*_*S*\*p*_ is the optimal monotonic classifier over *S* \ *p* in terms of *L*1−optimality.

For all parts *p* in *P*, the *L*_1_ of *C*_*S*\*p*_ over *p* is equal to the *L*_1_ of *C*_*S*\*p*_ over *S* minus the *L*_1_ of *C*_*S*\*p*_ over *S* \ *p*. According to Item 1, this *L*_1_ is greater than or equal to the *L*_1_ of *C*_*S*_ over *S* minus the *L*_1_ of *C*_*S*\*p*_ over *S* \ *p*. Furthermore, according to Item 2, this is also greater than or equal to the *L*_1_ of *C*_*S*_ over *S* minus the *L*_1_ of *C*_*S*_ over *S* \ *p*, which is equal to the *L*_1_ of *C*_*S*_ over *p*.

Knowing that, for all parts *p* of *P*, the *L*_1_ of *C*_*S*\*p*_ over *p* is greater than or equal to the *L*_1_ of *C*_*S*_ over *p*, by summing on all *p*, it holds that the sum of the *L*_1_ of *C*_*S*\*p*_ over all parts *p* is greater than or equal to the sum of the *L*_1_ of *C*_*S*_ over all partitions *p*, which is equal to the *L*_1_ of *C*_*S*_ over *P*. Therefore, it is greater than or equal to *L*_1_ of *C*_*S*_ over *S*. ?

And ∑*p*∈*P E*(*C*_*S*\*p*_, *p*) is the classification error using cross-validation over *P*. Therefore, MAE_CV_ ≥ MAE_full_.

### 4.2. Preselection Algorithm

The property above allows efficient pair detection solely based on MAE_full_, eliminating pairs that are bad due to a high MAE_full_. The preselection algorithm to identify the top pairs was based on this property. It identified pairs with favorable MAE_CV_. To achieve this, MAE_full_ was initially calculated for all pairs. Starting with the pairs having the lowest MAE_full_, their MAE_CV_ was computed. This process continued until the desired number of disjoint pairs—, i.e. pairs that do not contain the same genes,—was selected. Pairs with MAE_CV_ exceeding the maximum MAE_full_ among the selected pairs were discarded. To refine the selection threshold, MAE_CV_ were iteratively calculated for the remaining pairs, updating the maximum threshold whenever a new set of at least the target number of disjoint genes among the pairs was formed. This algorithm took as a parameter the number of disjoint genes which enable one to calibrate the outcome pairs.

The code for constructing MBMC and the preselection algorithm is available at https://github.com/oceanefrqt/MBMC

## 5. Discussion

This paper presents an extension of the monotonic bivariate classifiers to the multi-class setting, enabling the prediction of ordinal phenotypic outcomes based on transcriptomic data. The MBMC method demonstrates performance comparable to other state-of-the-art methods in predicting ordinal phenotypes, such as PFI and overall survival. However, one of the key advantages of MBMC is its nearly direct interpretability, as it identifies pairs of gene expressions that are associated with the phenotype of interest. It can enable the discovery of potential biomarkers and provide hypotheses about the mechanisms underlying the variability in outcomes.

Although we used pathway enrichment analysis to analyze the biological significance of the identified gene expression pairs, further investigation could be performed to understand the underlying biology. For example, the resulting gene expression pairs can be represented as a network of edges that connect gene expression pairs, which may provide further, higher-order insights. Advanced network analysis techniques can be applied to uncover patterns and biological signals within this network. In addition, future studies can explore the possibility of stratifying patients according to the gene expression pairs that are the most predictive of outcomes, potentially leading to more personalized and effective treatment strategies.

## Acknowledgments

OF, DA, KZ, SH, and BS have received funding from the European Union’s Horizon 2020 research and innovation programme under grant agreement No. 965193 for DECIDER. DA gratefully acknowledges support from Orion Research Foundation sr. CD acknowledges funding by the European Union (ERC, “dynaBBO”, grant no. 101125586). Views and opinions expressed are, however, those of the author(s) only and do not necessarily reflect those of the European Union or the European Research Council Executive Agency. Neither the European Union nor the granting authority can be held responsible for them.

## Footnotes

1 For reasons of brevity, from now on, we are referring to pairs of genes instead of pairs of gene expressions.

2 Note that these are the models obtained when trained on all the data, but the MAE-CVE corresponds to the MAE calculated with the 5-fold CV.

## References

J. S. Cardoso and R. Sousa. Measuring the performance of ordinal classification. International Journal of Pattern Recognition and Artificial Intelligence, 25(08):1173–1195, 2011. doi: 10.1142/S0218001411009093. URL https://doi.org/10.1142/S0218001411009093.

C.-K. Chen. The classification of cancer stage microarray data. Computer methods and programs in biomedicine, 108, 08 2012. doi: 10.1016/j.cmpb.2012.07.001.

O. Fourquet, M. S. Krejca, C. Doerr, and B. Schwikowski. Towards the genome-scale discovery of bivariate monotonic classifiers. bioRxiv, 2024. doi: 10.1101/2023.02.22.529510. URL https://www.biorxiv.org/content/early/2024/08/02/2023.02.22.529510.

E. Frank and M. Hall. A simple approach to ordinal classification. Lecture Notes in Computer Science, 2167:145–156, 08 2001. doi: 10.1007/3-540-44795-4_13.

L. Gaudette and N. Japkowicz. Evaluation methods for ordinal classification. In Y. Gao and N. Japkowicz, editors, Advances in Artificial Intelligence, pages 207–210, Berlin, Heidelberg, 2009. Springer Berlin Heidelberg. ISBN 978-3-642-01818-3.

P.A. Gutiérrez and S. García. Current prospects on ordinal and monotonic classification. Progress in Artificial Intelligence, 5, 03 2016. doi: 10.1007/s13748-016-0088-y.

A. Hakkinen, K. Zhang, A. Alkodsi, N. Andersson, E. P. Erkan, J. Dai, K. Kaipio, T. Lamminen, N. Mansuri, K. Huhtinen, A. Vaharautio, O. Carpén, J. Hynninen, S. Hietanen, R. Lehtonen, and S. Hautaniemi. PRISM: recovering cell-type-specific expression profiles from individual composite RNA-seq samples. Bioinformatics, 37(18):2882–2888, 03 2021. ISSN 1367-4803. doi: 10.1093/bioinformatics/btab178. URL https://doi.org/10.1093/bioinformatics/btab178.

D. C. Howell. Median Absolute Deviation, 2005. URL 10.1002/0470013192.bsa384.

A. Lahtinen, K. Lavikka, A. Virtanen, Y. Li, S. Jamalzadeh, A. Skorda, A. R. Lauridsen, K. Zhang, G. Marchi, V.-M. Identification of Monotonically Classifying Pairs of Genes for Ordinal Disease Outcomes

Isoviita, V. Ariotta, O. Lehtonen, T. A. Muranen, K. Huhtinen, O. Carpén, S. Hietanen, W. Senkowski, T. Kallunki, A. Häkkinen, J. Hynninen, J. Oikkonen, and S. Hautaniemi. Evolutionary states and trajectories characterized by distinct pathways stratify patients with ovarian high grade serous carcinoma. Cancer Cell, 41(6):1103–1117.e12, 2023. ISSN 1535-6108. doi: 10.1016/j.ccell.2023.04.017. URL https://www.sciencedirect.com/science/article/pii/S1535610823001435.

M. Luyckx, J.-L. Squifflet, A. M. Bruger, and J.-F. Baurain. Recurrent high grade serous ovarian cancer management. In S. Lele, editor, Ovarian Cancer. Exon Publications, 2022. ISBN 978-0-6453320-8-7. doi: 10.36255/exon-publications-ovarian-cancer-management. URL https://doi.org/10.36255/exon-publications-ovarian-cancer-management.

I. Nikolayeva, P. Bost, I. Casademont, V. Duong, F. Koeth, M. Prot, U. Czerwinska, S. Ly, K. Bleakley, T. Cantaert, P. Dussart, P. Buchy, E. Simon-Lorière, A. Sakuntabhai, and B. Schwikowski. A blood RNA signature detecting severe disease in young dengue patients at hospital arrival. The Journal of Infectious Diseases, 217(11):1690–1698, 2018. doi: 10.1093/infdis/jiy086.

F. Pedregosa, G. Varoquaux, A. Gramfort, V. Michel, B. Thirion, O. Grisel, M. Blondel, P. Prettenhofer, R. Weiss, V. Dubourg, J. Vanderplas, A. Passos, D. Cournapeau, M. Brucher, M. Perrot, and E. Duchesnay. Scikit-learn: Machine learning in Python. Journal of Machine Learning Research, 12:2825–2830, 2011.

G. Reifenberger, R. G. Weber, V. Riehmer, K. Kaulich, E. Willscher, H. Wirth, J. Gietzelt, B. Hentschel, M. Westphal, M. Simon, G. Schackert, J. Schramm, J. Matschke, M. C. Sabel, D. Gramatzki, J. Felsberg, C. Hartmann, J. P. Steinbach, U. Schlegel, W. Wick, B. Radlwimmer, T. Pietsch, J. C. Tonn, A. von Deimling, H. Binder, M. Weller, and M. Loeffler. Molecular characterization of long-term survivors of glioblastoma using genome- and transcriptome-wide profiling. International Journal of Cancer, 135(8):1822–1831, Oct 2014. doi: 10.1002/ijc.28836.

F. Stout. Isotonic regression via partitioning. Algorithmica, 66 (1):93–112, 2013. doi: 10.1007/s00453-012-9628-4.

Stroggilos, M. Frantzi, J. Zoidakis, M. Mokou, N. Moulavasilis, E. Mavrogeorgis, A. Melidi, M. Makridakis, K. Stravodimos, M. G. Roubelakis, H. Mischak, and A. Vlahou. Gene expression monotonicity across bladder cancer stages informs on the molecular pathogenesis and identifies a prognostic eight-gene signature. Cancers, 14(10), 2022. ISSN 2072-6694. doi: 10.3390/cancers14102542. URL https://www.mdpi.com/2072-6694/14/10/2542.

S. Tian. Identification of monotonically differentially expressed genes for non-small cell lung cancer. BMC Bioinformatics, 20, 4 2019. ISSN 14712105. doi: 10.1186/s12859-019-2775-8.

H. W. Wang, H. J. Sun, T. Y. Chang, H. H. Lo, W. C. Cheng, G. C. Tseng, C. T. Lin, S. J. Chang, N. R. Pal, and I. F. Chung. Discovering monotonic stemness marker genes from time-series stem cell microarray data. BMC Genomics, 16, 1 2015. ISSN 14712164. doi: 10.1186/1471-2164-16-S2-S2.

G. P. Zhuoqing Fang, Xinyuan Liu. Gseapy: a comprehensive package for performing gene set enrichment analysis in python. Bioinformatics, 2022. doi: 10.1093/bioinformatics/btac757.

